# scMinerva: an Unsupervised Graph Learning Framework with Label-efficient Fine-tuning for Single-cell Multi-omics Integrated Analysis

**DOI:** 10.1101/2022.05.28.493838

**Authors:** Tingyang Yu, Yongshuo Zong, Yixuan Wang, Xuesong Wang, Yu Li

## Abstract

Single-cell multi-omics is a rapidly growing field in biomedicine, where multiple biological contents, such as the epigenome, genome, and transcriptome, can be measured simultaneously. Despite its potential, the integrated analysis and prediction of cellular states based on this complex multi-omics data pose significant challenges due to data sparsity, high noise, and computational overhead. To address these challenges, we developed *scMinerva*, an unsupervised framework for single-cell multi-omics integrated analysis. The learned embeddings from the multi-omics data enable accurate integrated classification of cell types and stages. Specifically, we construct a heterogeneous graph from multiple omics and propose a novel biased random walk algorithm *omics2vec*, which can learn the heterogeneous biological graph in a way that balances both local and global network structures. scMinerva successfully outperforms existing unsupervised methods on various simulated and real-world datasets when fine-tuned by very few labels. Additionally, scMinerva demonstrates strong label efficiency, is robust to fluctuation in data quality, allows one omics to compensate for weakness in others and could effectively classify cells with different annotation granularities. Furthermore, we showcase scMinerva’s ability to accurately provide prospective biomarkers and predict cell differentiation trends for COVID-19-infected cells, through the joint analysis of multi-omics data.

## 1 Background

Single-cell technologies have revolutionized our understanding of biological systems by revealing the significant heterogeneity across different cell types and states. The simultaneous measurement of multiple omics, such as the epigenome, genome, and transcriptome, represents an exciting frontier for single-cell analysis [1, 2, 3]. Consequently, integrated analysis methods are required to handle complex multi-omics data for different downstream analyses. This is crucial for advancing the understanding of disease pathogenesis, identifying new therapeutic targets, and developing personalized treatments [4]. Despite the benefits, the inherent characteristics of the multi-omics data make it challenging to analyze computationally. One major challenge is the sparsity of single-cell data, which is due to the lack of gene activity in specific cell types and the relatively shallow sequencing of some droplet-based technologies [5]. Additionally, the high-dimensional features produced by the multiplexing and high throughput of multi-omics data can result in expensive computational overhead, making the analysis computationally expensive. Furthermore, the availability of single-cell annotations is limited as it is labor-intensive and requires expert knowledge, and inadequate samples could be given for some rare cases or conditions, making it difficult to study. These challenges have motivated researchers to develop various computational tools.

Current single-cell multi-omics tools can be broadly classified into three categories: 1) Latent-space inference methods, which assume a common latent space or kernel shared by all the omics and optimize it with matrix factorization [6, 7, 8] or manifold alignment [8, 9]. The utility of these tools can be limited by the strong assumption and they often require more sophisticated data preprocessing, 2) Correlation-based methods, which focus on the (dis)similarity measures that correlate different components to each other, including Seurat 4.0 [10], CiteFuse [11], Conos [12], scREG [13], etc. Their performance can degrade greatly when some omics are relatively noisy. 3) Deep learning-based methods, which train a model in an end-to-end manner, such as the recent unsupervised methods TotalVI [9] and DeepMAPS [14]. These methods can handle complex relationships between the different omics and are usually more robust to noise, but they mostly are designed to process CITE-seq data and therefore cannot handle datasets with more than two omics.

To summarize, existing methods still have several limitations: 1) Inability to process datasets with more than two omics (such as triple-omics), e.g., Seurat 4.0 [10], TotalVI [9] and DeepMAPS [14]. Forcibly integrating other omics will potentially impair performance. 2) Sensitivity to noise, especially the statistical methods. The performance of some existing methods may decrease when there is high noise in certain omics, which requires careful data pre-processing or may lead to failure analysis, e.g., MOFA+ [7]. 3) Challenging for adapting to annotation versions of fine granularity. Previously, researchers would typically employ unsupervised methods to classify sample cells by grouping them into different clusters. Then, they would use the aggregated cluster-level expression profiles and marker genes to label each cluster. However, this approach can be time-consuming and requires expert knowledge, particularly for cell annotations with finer granularity. 4) Lack of interpretability. Many existing methods, especially deep learning-based methods, provide little insight into the biological mechanisms underlying the integrated analysis results. This limits the ability of researchers to understand the biological significance of the findings and generate testable hypotheses.

To address these limitations, we propose **scMinerva**, an unsupervised and interpretable method that can flexibly handle any number of omics. It proposes a novel random walk strategy called “omics2vec” and leverages a graph convolutional network (GCN) model to jointly integrate the information. Specifically, our approach formulates the integration problem as a graph learning problem and constructs a heterogeneous graph by viewing each sample cell as a node and building sub-graphs for each omics. The model then learns a unified embedding for each cell by learning on this constructed graph in an unsupervised manner. Further, we introduce a new random walk strategy, omics2vec, that learns to embed nodes or samples by defining a biased random walk procedure. It runs random walk algorithm on the heterogeneous graph by distinguishing “breath-first search”, “depth-first search” and “omics-first search” node transition with biased transition probability. Therefore, this strategy explores the graph nodes’ similarity in a way that balances the exploration of local and global network structures, making it well-suited for the biological setting of single-cell integrated analysis.

With the output embeddings for sample cells from scMinerva, we focus on the single-cell classification tasks. Recent cell-type annotation methods for single-cell scRNA-seq suggest that, instead of aggregating cluster level expression profiles to classify sample cells, we can train some classifier by a small number of labeled cells (e.g. 10-20% randomly chosen from the sample pool) [15, 16]. Inspired by this, to effectively approximate and benefit the real-life classification tasks, we fine-tune a separate nearest neighbor classifier with very few labels (*i.e.*, 5%-20% of the data) on produced embeddings from scMinerva to predict cell types or stages. Moreover, since this classifier is independent of the unsupervised training stage, our embeddings can be fastly adapted to annotations with different granularity.

We extensively evaluate scMinerva on 4 simulated datasets and 6 real-world datasets, comparing it with previous state-of-the-art methods from different categories using 4 different metrics. Our method outperforms previous state-of-the-art methods by an average of ~15% on dataset SNARE, over 20% on GSE128639, and over 30% on simulated quadruple-omics datasets. Furthermore, compared to other methods, our approach is less sensitive to the amount of annotated data and has a stronger anti-noise ability when facing noisy omics. By comparing our results with the performance of single-omics baselines, we find that integrated analysis by scMinerva is robust and always better than any single-omics on classification tasks. We showcase the analysis of biomarker detection for Peripheral Blood Mononuclear Cells (PBMCs) using scMinerva. We demonstrate that scMinerva can identify meaningful biomarkers and that the gene activity value on these biomarkers can expressively reveal the cell stage in a fine-grained manner. More interestingly, we found that after training with GCN, nodes with a higher occurrence frequency have a higher probability to upregulate the marker genes compared to the counterparts before GCN training. This interprets the model in a way that if we assume nodes that have a higher chance to be walked as have a higher priority, GCN will assign a higher priority to high expression level nodes and more frequently utilize information from its neighbor. Furthermore, we use the predicted results of scMinerva to analyze the cell differentiation trend at the single-cell level for COVID-19-infected T cells.

We find that, for COVID-19 infected T cells, their differentiation to CD4^+^ T_CM_, CD4^+^ T_FH_, CD4^+^ prolif are activated. In summary, our contributions are as follows:

- We introduce an unsupervised and interpretable method scMinerva by constructing a heterogeneous graph from multi-omics data, propose a new biased random walk algorithm omcis2vec to this biological setting, and then learn unified embedding for each sample cell.
- Our method outperforms previous state-of-the-art methods on integrated cell-type classification and can integrate any number of omics types. We extensively evaluate its label efficiency, antinoise ability when facing noisy omics, and robustness that can benefit from different single omics. Moreover, it allows for faster adaptation of the embeddings to annotations with varying levels of granularity.
- To prove its practical utility, we showcase the analysis of biomarker detection and single-cell differentiation trends using the results from scMinerva. With the help of scMinerva, researchers can effectively identify meaningful biomarkers, analyze single-cell differentiation, *etc.*

## 2 Results

### 2.1 ScMinerva is designed for single-cell multi-omics integrated analysis

In this section, we introduce a novel method called *scMinerva* for unsupervised single-cell multi-omics integrated analysis as illustrated in Figure 1. Single-cell multi-omics data contains gene activity values for each cell from various omics, each of which measures different biological processes. The goal of integrated analysis is to generate comprehensive embedding for each sample cell from the multi-omics data. To achieve this, scMinerva formulates the problem as a unified heterogeneous graph learning task. In the framework of scMinerva, we employ a graph convolutional network (GCN) on heterogeneous graphs and proposes a biased random walk algorithm called *omics2vec* with two variances “omics2vec-for-nodes” and “omics2vec-for-samples”.

**Figure 1:**
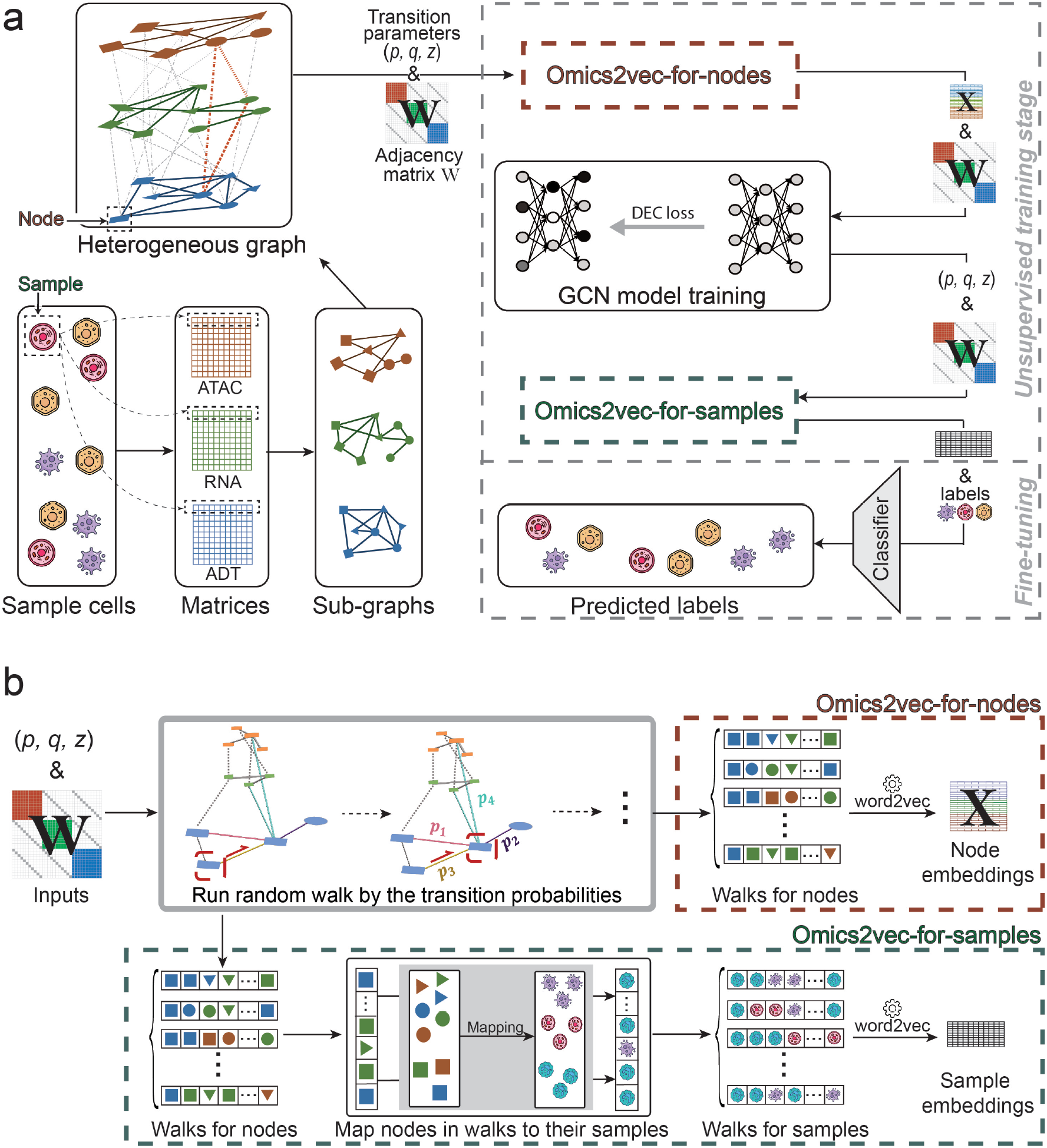
ScMinerva workflow and the omics2vec algorithm. **a.** ScMinerva takes the preprocessed single-cell gene activity matrices as inputs, and performs integrated analysis on a constructed heterogeneous graph. In the framework, we propose an algorithm called omics2vec and leverage the graph convolutional network (GCN) model to learn unified sample embeddings. **b.** Omics2vec is a graph learning algorithm based on biased random walk that offers two versions. Both versions rely on the transition parameters (*p, q, z*) and the adjacency matrix **W** as inputs, and apply a biased random walk to explore the graph. The first version, omics2vec-for-nodes, aims to embed the nodes of the heterogeneous graph. This variant directly takes the walks generated for nodes and feeds them into the word2vec model, which then produces the node embeddings. On the other hand, omics2vec-for-samples focuses on embedding the samples and completing integrative analyses. This variant maps the indices of nodes in the walks to the indices of samples, and then feeds the resulting walks for samples into the word2vec model, which generates sample embeddings.

Omics2vec is inspired by word2vec [17] and the node2vec algorithm [18]. Word2vec is a model that learns to embed words in a high-dimensional space, where semantically similar words are grouped together. Node2vec extends this idea to graph-structured data by defining a biased random walk procedure that explores the similarity of nodes in a way that balances the exploration of local and global network structures. This algorithm combines breadth-first search (BFS) and depth-first search (DFS) techniques and inputs the generated walks into word2vec to ensure that the embeddings of spatially close nodes are similar. However, applying node2vec to a biological heterogeneous graph presents a challenge because the inter-omics transition edges are undefined. Additionally, the weights of the inter-omics edges play a significant role in the performance. They determine the probability of being explored in the neighborhoods of a sample’s counterparts on different omics. To overcome this challenge, we propose a novel algorithm called omics2vec, which has two variances: “omics2vec-for-nodes” and “omics2vec-for-samples”. These variations extend the node2vec idea to analyze biological heterogeneous graphs.

Our method comprises four essential steps to facilitate a comprehensive analysis of multi-omics data. Firstly, we create a heterogeneous graph from multi-omics data, allowing us to achieve a unified learning approach. Secondly, we run “omics2vec-for-nodes” to learn node embeddings on the heterogeneous graph. This approach maps the nodes in the graph to a low-dimensional space, where similar nodes are located closer to each other. This technique enables us to capture the intrinsic relationships among the different omics types in a more comprehensive manner. Thirdly, we train a Graph Convolutional Network (GCN) model to gather similar nodes and disperse dissimilar nodes on the heterogeneous graph. The GCN model aggregates information from the neighboring nodes and enables us to obtain a more accurate representation of the relationships among local nodes. Finally, we employ “omics2vec-for-samples” to learn sample embeddings from the trained heterogeneous graph. The output sample embeddings allow us to analyze the sample cells in an integrated view. Now, we will discuss each step in detail.

The first step in scMinerva is to construct a heterogeneous graph. We separately build weighted K-nearest neighbor (KNN) graphs for each omics, which we refer to as sub-graphs. For each sample cell, there is a mapping node on each sub-graph, which may hold complementary information. To learn all the information from different omics, we link the mapping nodes of a sample cell in one sub-graph with its corresponding mapping nodes in the other sub-graphs. This step creates a heterogeneous graph that contains similarity information from all omics and enables us to learn a unified embedding for sample cells.

In the second step of scMinerva, we generate embeddings for nodes on the heterogeneous graph using the “omics2vec-for-nodes” algorithm. Instead of the mixture of BFS and DFS used by node2vec, we guide the biased random walk with three parameters: *p, q*, and *z*, all of which are positive values. The parameter *p* controls the probability for BFS, *q* controls DFS, and *z* is a new addition to the algorithm that controls the “omics-first search”. These hyper-parameters define the transition probability of the random walk procedure from one node to another. We further investigate the parameter sensitivity of this algorithm in Appendix. We can generate random walks on the graph controlled by these hyper-parameters, input the generated walks into word2vec, and obtain the embeddings of nodes.

During the third step, we utilized graph convolutional network (GCN) to optimize the node embeddings obtained in the previous step. We fed both the embeddings and the topology of the heterogeneous graph into the GCN model and trained it using an unsupervised loss function. This loss function aimed to disperse dis-similar nodes on the heterogeneous graph while gathering similar nodes, ultimately improving the quality of the embeddings. Specifically, we followed the work of Caron et al. [19] and used their unsupervised loss function DeepCluster, which encourages the embeddings of connected nodes to be close in the embedding space while pushing away the embeddings of disconnected nodes. The detailed description of the GCN training process can be found in the Method section.

In the fourth step of the scMinerva algorithm, we use another variance of omics2vec called “omics2vec-for-samples” to generate unified sample embeddings. While “omics2vec-for-nodes” generates node embeddings for each node on the graph, our goal is to obtain embeddings for individual sample cells. To achieve this, we remap nodes’ indices to their respective samples’ indices. Specifically, we ensure that the counterparts of one sample cell from different omics are mapped to the same sample cell in the walks. By inputting these remapped walks into the word2vec model, we obtain the embeddings of sample cells that leverage neighborhood information from the entire heterogeneous graph. This approach effectively captures integrated multi-omics information and completes the unsupervised training stage of the scMinerva algorithm. With the output sample embedding from scMinerva, we independently fine-tune the model with a nearest-neighbor classifier to fit the sample embeddings and predict their cell types or stages.

### 2.2 scMinerva outperforms previous state-of-the-art methods in single-cell integrated classification

In this section, we provide a comprehensive evaluation of our method’s performance on both simulated and real-world datasets. Specifically, we assess the anti-noise ability of our method using simulated datasets in Section 2.2.1, and its accuracy using real-world datasets in Section 2.2.2. We acknowledge that in real-life scenarios, cell-type annotation tasks are often completed by clustering cells first and using aggregated cluster-level expression profiles and marker genes to label each cluster. However, this approach can be challenging for fine-grained cell-type/state annotations and requires expert knowledge. To facilitate this process, we evaluated our method in a label-efficient manner using a simple nearest-neighbor classifier. This classifier is independent of the unsupervised training stage and is able to adapt the embeddings to annotations with different granularity. Specifically, we fit the produced embeddings of scMinerva and other methods using independent nearest neighbor-based classifiers and fine-tuned them using only 10% randomly chosen annotations from the entire training set. This approach eliminates the performance dependence on the results of clustering and allows for faster adaptation of the embeddings to annotations with varying levels of granularity. For additional details on the experimental settings, please refer to Section 4.2.6.

#### 2.2.1 Simulated datasets

The increasing trend in the development of experimental methods that jointly profile three or more omics types is widely recognized, for example, the combination of single-cell RNA sequencing (scRNA-seq), single-cell chromatin accessibility profiling (scATAC-seq), single-cell methylome profiling (scMethyl-seq), and single-cell proteomics [20]. However, as the number of omics types increases, it becomes harder to identify common latent features or sample correlations. For instance, handling triplet-omics datasets is more challenging than couplet-omics datasets due to the different distributions of gene activity matrices and potential pollution of certain omics by natural or artifact noises. Therefore, there is a pressing need for computational algorithms with stronger anti-noise capabilities to handle more omics data, including triplet-omics and quadruplet-omics datasets. To address this, we first simulate pseudo-multi-omics datasets to explore the impact of noise on the performance of integrated analysis methods as the number of omics types increases.

Since there exist real-world couplet-omics or triplet-omics datasets, we leave the evaluation on real-world datasets with two omics or triplet-omics to the next subsection. To evaluate the anti-noise ability of methods while the omics type number increases, we generated simulated quadruplet-omics datasets with four pseudo-omics. Our goal was to produce simulated datasets that contain different simulated gene activity matrices with varying distributions and certain noisy omics. To accomplish this, we used the single-cell RNA (scRNA) data simulator, splatter, and three convolutional neural networks (CNNs) [21]. We obtained simulated RNA-seq by inputting the RNA-seq data from GSE156478-CITE [22]. We trained the CNNs with real-world datasets to learn commonly shared latent features for omics. Specifically, we trained the networks by mapping sci-CAR RNA-seq to its ATAC-seq, GSE156478-CITE RNA-seq to its ADT data, and GSE156478-ASAP ATAC-seq to its ADT. By inputting the simulated RNA-seq to the trained networks, we generated four-omics datasets with 5 classes and sample numbers 2k, 5k, 10k, and 30k, respectively. These simulated datasets contained pseudo-RNA-seq (PO1 shown in Table 1), pseudo-ATAC-seq (PO2), pseudo-ADT from RNA-seq (PO3), and pseudo-ADT from ATAC-seq data (PO4). We acknowledge that simulating quadruplet-omics datasets can be a challenging task. To ensure that our simulated datasets were not trivial linear regression tasks, we assumed the presence of commonly shared latent features for real-life omics and trained the CNNs to learn these features. Further details on data simulation can be found in Data simulation from real-world datasets.

**Table 1:** Performance comparison of several methods on simulated quadruplet-omics data. The first column of the table presents the number of samples in each dataset. In the table, the best-performing method is denoted by bold text, while the second-best method is underlined. Columns labeled PO1 to PO4 represent the performance of four different single-omics methods. For each method, we used a KNN classifier and took 10% of the available ground-truth labels as the training set. Our results demonstrate that scMinerva consistently outperformed the second-best method across all metrics, achieving a significant improvement of approximately 30%. These results confirm the effectiveness of scMinerva in integrating different types of omics data for improved performance in downstream analyses.

| Size | Metric | scMinerva | MOFA+ | Conos | Seurat 4.0 | PO1 | PO2 | PO3 | PO4 |
| --- | --- | --- | --- | --- | --- | --- | --- | --- | --- |
| 2k | ACC | **0.911** | 0.474 | <u>0.627</u> | 0.263 | 0.905 | 0.229 | 0.351 | 0.217 |
|  | F1-weighted | **0.913** | 0.464 | <u>0.616</u> | 0.209 | 0.906 | 0.199 | 0.333 | 0.182 |
|  | F1-macro | **0.913** | 0.462 | <u>0.615</u> | 0.211 | 0.907 | 0.199 | 0.332 | 0.182 |
|  | ARI | **0.781** | 0.127 | <u>0.307</u> | 0.004 | 0.775 | 0.005 | 0.060 | 0.002 |
| 5k | ACC | **0.952** | 0.547 | <u>0.764</u> | 0.411 | 0.976 | 0.236 | 0.536 | 0.239 |
|  | F1-weighted | **0.952** | 0.548 | <u>0.764</u> | 0.408 | 0.976 | 0.223 | 0.524 | 0.224 |
|  | F1-macro | **0.952** | 0.548 | <u>0.656</u> | 0.407 | 0.976 | 0.224 | 0.522 | 0.224 |
|  | ARI | **0.913** | 0.198 | <u>0.449</u> | 0.099 | 0.942 | 0.007 | 0.250 | 0.009 |
| 10k | ACC | **0.957** | 0.691 | <u>0.719</u> | 0.220 | 0.986 | 0.262 | 0.399 | 0.211 |
|  | F1-weighted | **0.957** | 0.686 | <u>0.714</u> | 0.165 | 0.986 | 0.258 | 0.389 | 0.206 |
|  | F1-macro | **0.957** | 0.686 | <u>0.714</u> | 0.164 | 0.986 | 0.258 | 0.388 | 0.206 |
|  | ARI | **0.895** | 0.410 | <u>0.450</u> | 0.001 | 0.966 | 0.017 | 0.087 | 0.000 |
| 30k | ACC | **0.968** | <u>0.675</u> | 0.447 | 0.256 | 0.996 | 0.275 | 0.300 | 0.265 |
|  | F1-weighted | **0.968** | <u>0.676</u> | 0.446 | 0.230 | 0.996 | 0.272 | 0.297 | 0.261 |
|  | F1-macro | **0.968** | <u>0.677</u> | 0.446 | 0.230 | 0.996 | 0.272 | 0.297 | 0.261 |
|  | ARI | **0.927** | <u>0.377</u> | 0.134 | 0.007 | 0.990 | 0.024 | 0.024 | 0.025 |

**Table 2:** Datasets analyzed in this paper.

| Dataset | Cell Number | Cell Type | #Omics | Depth | Dropout |
| --- | --- | --- | --- | --- | --- |
| GSE128639 | 30k | PBMC | 3 | 11k | 95% |
| GSE156478-CITE | 4.6k | PBMC | 2 | 6k | 92% |
| GSE156478-ASAP | 4.5k | PBMC | 2 | 5k | 96% |
| COVID-PBMC | 79k | PBMC | 3 | 100k | 81% |
| scNMT | 875 | Mouse Embryonic Stem Cells | 3 | 25k | 77% |
| SNARE | 1047 | Mouse Cerebral Cortices | 2 | 9.5k | 92% |

In order to evaluate the performance of the different methods, we fine-tuned an 8-nearest neighbor classifier with 10% label on the embeddings outputted by all the methods. The results of our analysis are presented in Table 1, where we compared the performance of our proposed method with Seurat 4.0 [10], MOFA+ [7], Conos [12], and using only single pseudo-omics. Notably, we did not include TotalVI [9], CiteFuse [11], and DeepMAPS [14] in this comparison as they are not capable of processing more than two omics. We plan to compare these methods in the later section that focuses on two omics real-world datasets. We evaluated the performance using four metrics, namely accuracy, F1-weighted score, F1-macro score, and adjusted rand score (ARI). The details on the evaluation metrics used can be found in Method Performance evaluation. Overall, our results demonstrate that the proposed method outperformed the other methods in terms of accuracy, F1-weighted score, and ARI across all sample sizes.

Our results in Table 1 demonstrate that scMinerva outperforms all other evaluated methods in terms of accuracy, F1-weighted score, F1-macro score, and adjusted rand score. Specifically, scMinerva shows a significant improvement of around 30% over the second-best method in all metrics and all simulated datasets. However, we note that scMinerva’s performance could not surpass that of the best-performing single pseudo-omics PO1. This is because the simulated datasets were generated without considering complementary biological information, and scMinerva’s performance heavily relies on the presence of biologically relevant information in the input data. We also highlight that most existing methods, particularly statistical methods, are highly sensitive to low-quality omics in the datasets and perform poorly. In contrast, scMinerva demonstrates stronger robustness against noisy and challenging omics, as evidenced by its stable performance across all simulated datasets. Overall, scMinerva stands out as a promising method for analyzing multi-omics datasets with anti-noise capabilities.

#### 2.2.2 Real-world datasets

The previous subsection has demonstrated that our method exhibits superior performance over existing techniques in challenging scenarios, such as quadruplet-omics datasets or datasets containing extremely noisy pseudo-omics. To further validate the efficacy of our approach, we conduct experiments on real-world datasets. Similar to the previous subsection, we perform integrated classification experiments in a label-efficient manner using a simple nearest-neighbor classifier. Specifically, we generate embeddings for scMinerva and other methods and fit them all with an independent nearest neighbor-based classifier that is fine-tuned using only 10% annotations from the entire training set.

We evaluate our approach on several datasets, including CITE-seq (GSE128639 [23], GSE156478-CITE [22], COVID-PBMC [24]), ASAP-seq (GSE156478-ASAP [22]), SNARE-seq [25], and scNMT-seq [26]. These datasets exhibit diverse sample sizes, ranging from 1k to 64k. For the COVID-PBMC dataset, we segregate healthy samples and critical symptom samples, forming the COVID-non-covid dataset and the COVID-critical dataset, respectively. We extract these two datasets for later analysis. All datasets undergo standard pre-processing and quality control, as detailed in the Method 4.1. Prior to pre-processing, most of these datasets have a dropout rate exceeding 90% and shallow measurement depths, as listed in Figure 3.

We conducted experiments using various existing methods, including DeepMAPS, CiteFuse, to-talVI, and Seurat 4.0. These methods were run using their default settings. For MOFA+ and Conos, we adjusted the dimension of their embeddings to 200 to ensure the retention of sufficient latent information, similar to other methods. However, it is important to note that MOFA+ was constrained by memory limitations. When the number of samples exceeded 10k, we failed to run the originally preprocessed data on MOFA+ with 32G RAM. As a solution, we applied PCA on the data with 300 components for dataset GSE128639. Additionally, DeepMAPS, CiteFuse, and TotalVI are unsuitable for cases with three omics, and therefore, they are not included in the chart for COVID-PBMC, scNMT, and GSE128639. We evaluated the classification performance using the generated embeddings by fitting a K-nearest Neighbor (KNN) Classifier. For datasets containing more than 5k samples, we set the number of neighbors to 30, and for datasets smaller than 5k, we set it to 8. The sklearn library [27] was used for this purpose. The details of the data preprocessing are listed in Method Datasets, annotations and pre-processing, while a comprehensive description of the experimental settings can be found in Section Method. The results of these experiments are presented in Figure 2.

**Figure 2:**
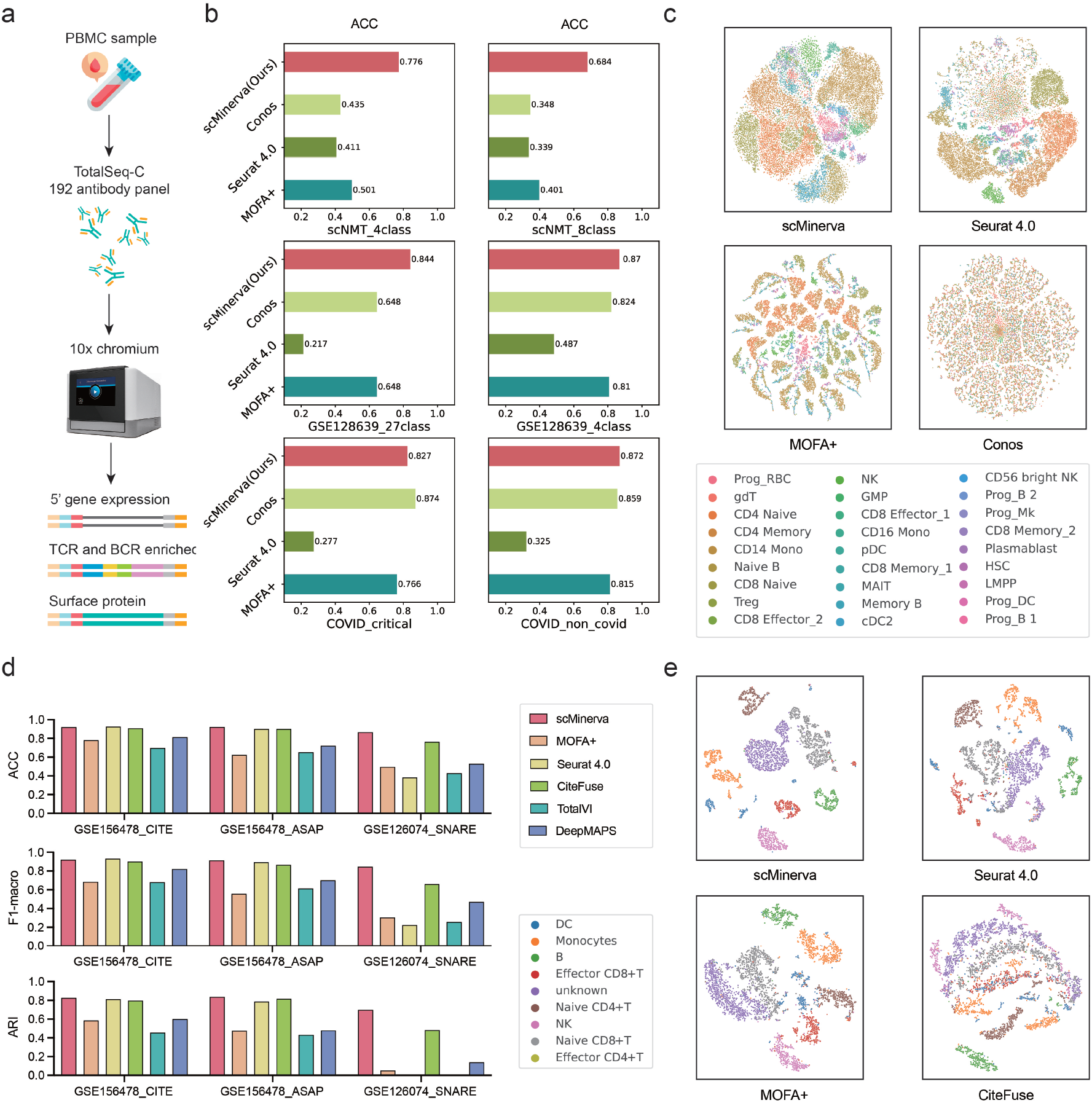
Classification performance on real-world datasets. **a**. The sequencing technique for Peripheral blood mononuclear cells to obtain three-omics data. **b**. Classification accuracy comparison on four triplet-omics datasets with six sets of annotations. We fine-tune the output embeddings with 10% labels. Our method shows a overall-the-best performance. **c**. Visualization on three-omics dataset GSE128639 with 30k cells and 27 classes for scMinerva, Conos, Seurat 4.0, and MOFA+. scMinerva’s visualization has the clearest boundary and separates clusters properly. **d**. The classification performance on two-omics datasets for scMineva and five exiting methods. We fine-tune the output embeddings with 10% labels. scMinerva always shows both great and stable ability even on SNARE which has one extremely noisy omics. **e**. Visualization on dataset GSE156478-ASAP with 5k cells and 8 classes. The scatter is colored by ground-truth labels. It is obvious that scMinerva nicely has the most dispersed clustering which best matches the ground-truth clusters.

With the exception of scNMT-seq, which measures the differentiation of mouse embryonic stem cells, most of the existing three omics datasets are based on human peripheral blood mononuclear cells (PBMC). In contrast to embryonic stem cells, where measurements typically include ATAC-seq, RNA-seq, and DNA methylation levels, PBMCs consist of lymphocytes (T cells, B cells, NK cells) and monocytes. Therefore, measurements of T cell receptor (TCR) and B cell receptor (BCR) are available in addition to rich expression levels of surface proteins, as shown in Figure 2a.

The algorithm scMinerva has been evaluated on several triplet-omics datasets, including scNMT, GSE128639, and COVID-PBMC, with a 10% fine-tuning data of different methods. The results are listed in Figure 2b, which shows that scMinerva outperforms most of the compared integrative analysis methods on classification tasks and has comparable performance with Conos, the best-performance method on the COVID_critical dataset. scMinerva exhibits notable improvements in classification accuracy, with an average increase of 7% and up to 20% on GSE128639 when classifying into 27 classes compared to the second-best method. It is important to note that scMinerva’s strength in classifying fine-grained cell classes is more evident. Given that the output of all methods is fine-tuned for classification using the same independent classifier, it is possible to compare their performance fairly under classification tasks with different annotation granularity. In the case of dataset GSE128639, scMinerva outperformed Conos by approximately 5% when classifying cells into four main cell types. However, when classifying sample cells into 27 sub-types, scMinerva showed a remarkable improvement of 20%. These results clearly demonstrate scMinerva’s capability in accurately classifying cells at a more granular level. Furthermore, scMinerva demonstrates its stability when only 10% labels are used for training. Importantly, different datasets have various versions of data annotations with different numbers of cell types. For example, GSE128639 is annotated by 4 classes and 27 sub-classes. The unsupervised integrated stage and an independent fine-tuning scheme of scMinerva ensure a faster adaptation to different data annotation granularity. To further evaluate the performance of scMinerva, we visualize the embeddings of the GSE128639 dataset produced by scMinerva, Conos, Seurat 4.0, and MOFA+ via t-SNE in Figure 2**c**. GSE128639 contains 30k cells across 27 cell types and cell states. From the scatter plot colored by the ground truth labels, it is evident that scMinerva clusters samples from the same type together and shows clear boundaries between different types. In contrast, MOFA+ shows a dispersed gathering for samples that should be in the same cluster. The embeddings of different cell types in the visualizations of Seurat 4.0 and Conos overlap, indicating that they are not able to effectively discriminate different cell types. These findings strongly support the good overall performance of scMinerva in the classification of triplet-omics datasets.

Moreover, couplet-omics PBMC datasets, GSE156478-CITE and GSE156478-ASAP, were used to test the performance of existing methods for classification. The results in Figure 2**d** show that scMinerva achieved a slightly better performance than the second-best method on these two datasets. Seurat 4.0 and CiteFuse also showed similar performance with scMinerva, but CiteFuse was not adaptive to datasets containing more than two omics, and Seurat 4.0 had poor performance on high-noise datasets such as SNARE. The embedding of GSE156478-ASAP was visualized in Figure 2**e**, and scMinerva was found to have a clearer embedding than the other three methods. However, on the SNARE dataset, scMinerva outperformed all existing methods on classification with an around 15% promotion. This performance gap occurred because scMinerva had excellent anti-noise ability and could effectively handle situations with a severe quality gap between different omics. The result listed below in Figure 3 shows a severe quality gap between two omics of the SNARE dataset, as the performance of ATAC-seq was very poor. However, scMinerva was not affected by the poor quality of certain omics and showed strong robustness. On the other hand, all other methods, including TotalVI, DeepMAPS, Seurat 4.0, and MOFA+, tended to mess up all samples to one class, while CiteFuse was strongly encumbered by the high noise in ATAC-seq. scMinerva achieved an accuracy of over 80% and had a 20% improvement on ARI over other methods.

**Figure 3:**
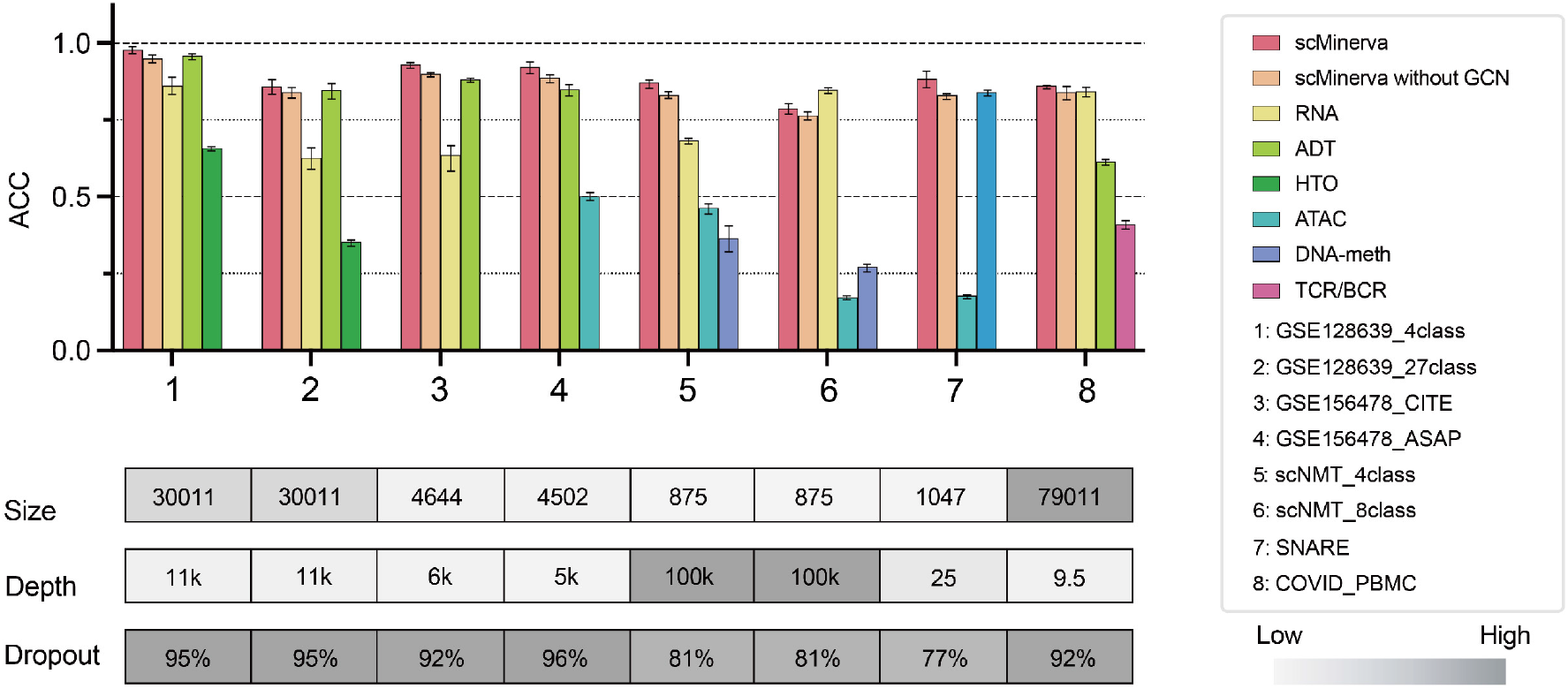
Ablation study results on real-world datasets. The clustered histograms represent results from the same datasets, and below them, we list their sample sizes, measurement depths, and dropout rates. The error bars are the standard deviation of 10 times runs with different random seeds. It shows that scMinerva achieves the best performance in most cases and efficiently integrates valid information from different omics. In some cases, scMinerva without GCN training will produce a lower performance than the best single omics data. However, with a GCN implemented, the performance mostly beats any single omics and is promoted to a higher level.

In conclusion, scMinerva has demonstrated strong overall performance in multi-omics data integration compared to existing methods. Its ability to effectively handle challenging scenarios with severe quality gaps between different omics and its robustness to noise make it a promising approach.

### 2.3 Integration on multi-omics data is robust and overall better than any single omics on classification

With good accuracy and anti-noise ability, we further evaluated whether scMinerva could efficiently capture valid information from different omics and obtains a more comprehensive inference. To achieve this, we perform an ablation study on real-world datasets. The performance of the proposed multi-omics integration method is compared with that of the single-omics data. For each omics, we separately build the graph topology, run the random walk without the omics-transition parameter, generate embeddings, and classify the embeddings with same nearest neighbor classifier fine-tuned by 10% data. To enable a fair comparison, we ran the experiments 10 times for each dataset. Moreover, the data split across all omics for each time is from fixed random seeds with the same proportion and hyper-parameters. Also, to validate the necessity of using GCN, we compared the results before and after training with all other components fixed.

The findings presented in Figure 3 demonstrate that combining multi-omics data using GCN yields better performance than using any single omics data in 8 out of 9 datasets. The only exception was scNMT for 8-way classification, where there was a mere 3% drop compared to the best-performance omics. This could be attributed to the homogeneity of cell states in the other two omics data, apart from RNA-seq. As a result, the graph topologies from ATAC-seq and DNA-meth were almost random under a KNN graph construction, potentially causing our method to be influenced when starting to walk from some extremely low-quality nodes. Furthermore, our proposed method, scMinerva with GCN, achieved an overall 2%-8% improvement compared to the model without GCN training. This is because the DEC loss guided GCN model to gather similar nodes in a global scenario for the heterogeneous graph, which made scMinerva more robust and enabled it to integrate global information jointly from all the omics data. In summary, our results indicate that scMinerva efficiently captures valid information from different omics and yields a more comprehensive inference in most cases.

### 2.4 scMinerva shows strong scalability on label efficiency

In addition to robustness and accuracy, the scalability of methods across different levels of labeled data is also an important practical consideration. To this end, we fine-tuned classifiers using only 5%, 10%, and 20% labeled data as the training set for all the methods. The performance of scalable and label-efficient methods should not suffer significantly as the size of the training set decreases. We evaluated the performance of the methods using four metrics: F1-weighted, F1-macro, ARI, and ACC, and applied them to four triplet-omics datasets. In order to ensure a rigorous and unbiased comparison, we conducted experiments on each dataset and method a total of 5 times. To obtain reliable and consistent results, we calculated the average performance across these 5 trials. Furthermore, we took measures to ensure that our testing process was standardized across all methods. Specifically, we used fixed random seeds to split the data for fine-tuning the classifiers in each test, with the same proportion utilized across all methods. By adopting these practices, we aimed to minimize the impact of potential confounding factors and obtain accurate and meaningful results.

As shown in Figure 4, we displayed the performance of methods which can handle triplet-omics. Each row contains three ticks that represent a method’s performance when fitted with classifiers with 5%, 10% and 20% labels from left to right, respectively. Thus, a more scalable and label-efficient method should exhibit shorter ticks. From Figure 4, it can be easily observed that for all the datasets and train-test splits except the COVID_critical dataset, our method not only has the best performance but also has relatively small variances compared with others. Especially for the COVID_non_covid dataset and GSE128639 dataset, the performances of scMinerva are not only the best but also the most robust. Its performances are nearly unchanged while the fine-tuning label size is dropped from 20% to 5%. For the COVID_critical dataset, Conos achieves the best performance while our method has comparable performances and smaller various compared to Conos. In the scNMT dataset, we used approximately 50 randomly-selected labeled cells, which represents 5% of the dataset, to refine the output of all the approaches. Despite this relatively small sample size, we were able to achieve an accuracy of 74.3% on 4-way classification. Notably, other existing methods that were tested proved unsuccessful, as they tended to misclassify all samples into a single class.In summary, scMinerva shows the overall best scalability compared to other methods. The power of scMinerva is sparkled by easy fine-tuning and is not sensitive to the using label size for fine-tuning a separate classifier. This label efficiency reveals great potential in practice for reducing the labor and resource consumption for single-cell data annotation.

**Figure 4:**
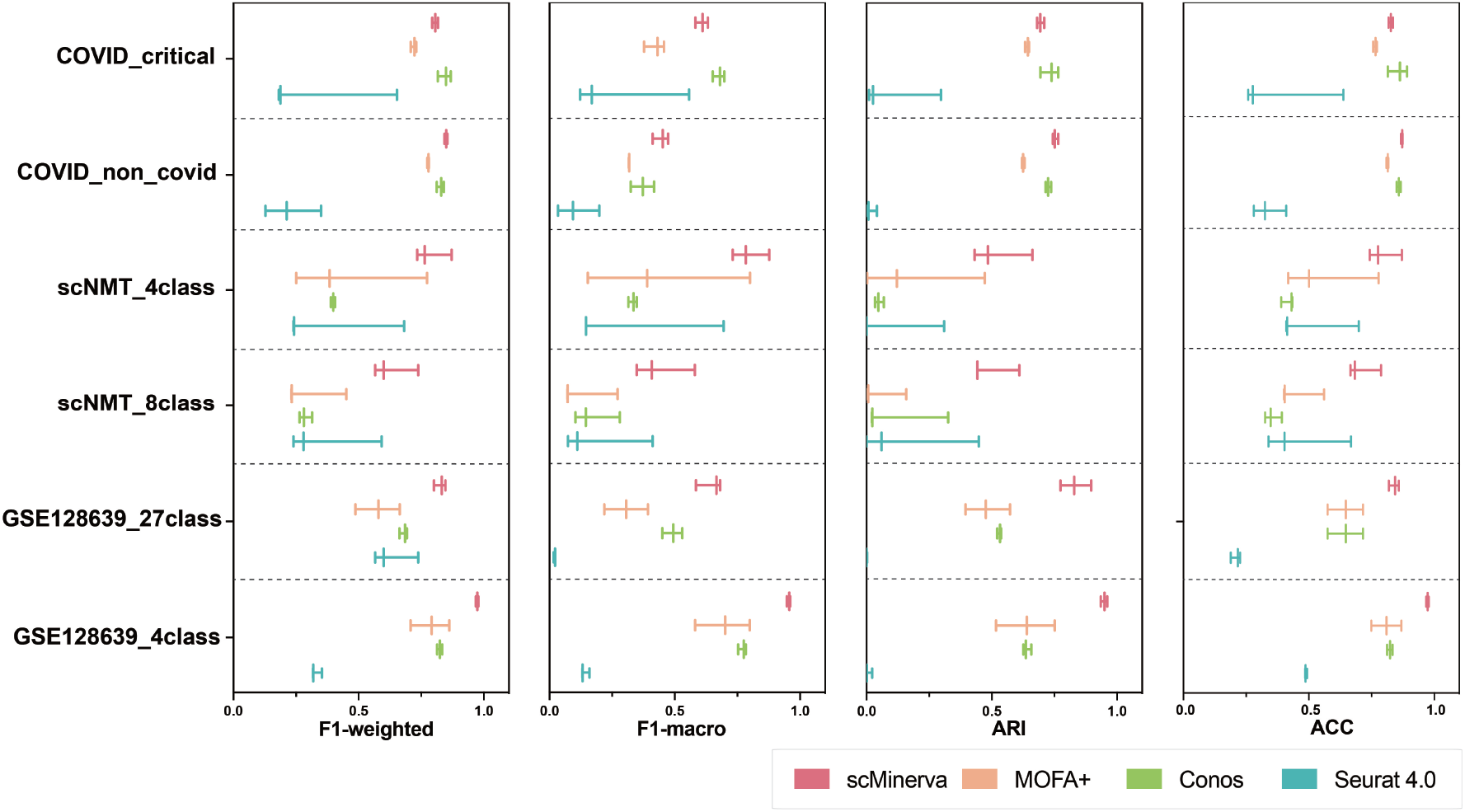
Scalability evaluation by fine-tuning classifiers with varying amounts of labeled data and measured the integrated classification performance. The results are presented in Figure X, where each row represents the performance of the classifiers fine-tuned on 5%, 10%, and 20% of the labeled data, respectively. The length of the ticks indicates the performance of each method. As shown in the figure, scMinerva achieves the best overall performance and demonstrates stronger scalability compared to other methods. Notably, our method exhibits shorter ticks, indicating better scalability, particularly in scenarios with limited labeled data.

### 2.5 scMinerva accurately identifies biomarkers of PBMC cells

To further demonstrate the versatility of scMinerva, we conducted integrated analysis on PBMC multi-omics datasets that contained 5 general cell types. Our aim was to identify meaningful biomarkers that could act as indicators of biological processes and play a vital role in disease detection [28]. We used scMinerva to predict cell types and detect biomarkers on the predicted clusters using SCANPY [29]. Our experiments were conducted on the GSE128639 dataset, which contained 5 general cell types and 27 subclasses across T cell, B cell, progenitor cell, NK cell, and Mono/DC cell.

We first identified genes that were highly expressed in the 5 general cell types, as shown in Figure 5**a**. The X-axis represented the rank of their expression level in each cell type. To further demonstrate that these detected genes were more highly expressed in the specific cell types than in the rest of the cell types, we selected the top 5 marker genes in each cell type and plotted violin graphs for each cell type to compare their expression levels, as shown in Figure 5**b**. The blue color represented the expression of the genes in that particular cell type, while the orange color represented the sum of the expression in the rest of the cell types. The results indicated that the expression level of the detected marker genes was significantly higher in the specific cell types than in the rest of the cell types. From a practical perspective, the detected biomarkers can reveal latent information on their relative biological processes. For example, the gene MALAT1, detected to be ranking 5 in B cells, is shown to be suitable to act as a biomarker, as it correlates with larger tumor size, advanced tumor stage and overall poor prognosis [30]. This evidence mutually confirms the effectiveness of the biomarker detection of scMinerva.

**Figure 5:**
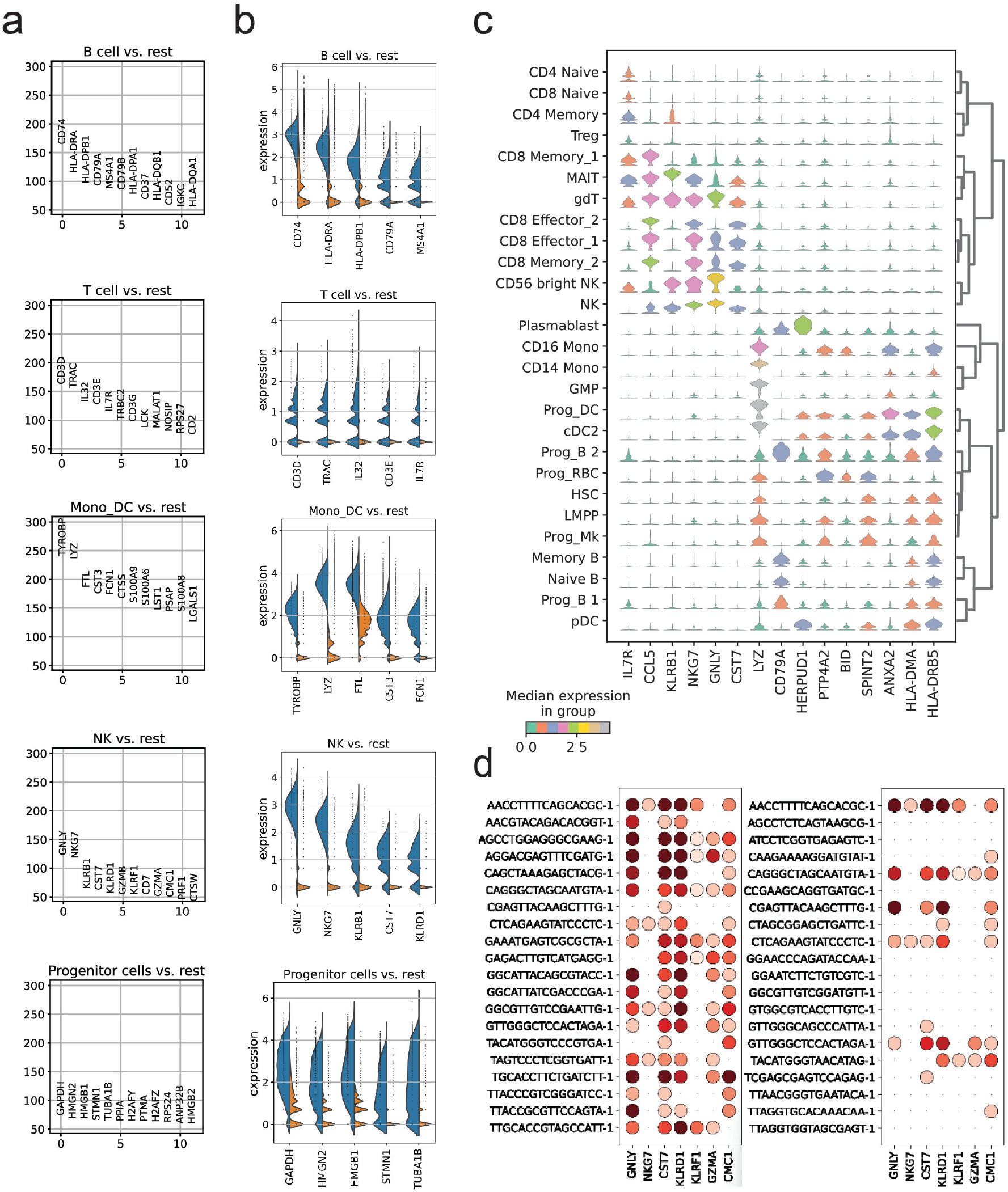
Biomarker detection was performed on PBMC cells from the GSE129639 dataset. **a.** The detected marker genes for 5 general cell types. **b.** The comparison of the expression level between the the top 5 detected biomarkers in the corresponding cell type and the rest of the cell types. **c.** A violin graph depicting the expression level of the biomarkers in the 27 subclasses. **d.** The expression levels of the nodes of most frequent occurrence in walks on marker genes before and after training. While there were no significant differences in expression levels between the 20 cells with the most frequent occurrence in walks before training, the top 20 nodes after training showed a strong upregulation of marker genes. These results suggest that the nodes with high feature expression levels are important in the generation of output embeddings, especially in sparse single-cell data lacking high library-size cells.

To further show that scMinerva’s predictions could be used to detect biomarkers in more fine-grained classes, we conducted experiments on the 27 subclasses. Figure 5**c** depicted the results, with the vertical axis representing the 27 subclasses and the horizontal axis indicating the detected marker genes. The color of the violin graphs represented the expression level of these genes in the corresponding cell types. For instance, gene LYZ was highly expressed in cell types GMP, Prog-DC, and cDC2, which was also confirmed by [31, 32]. The detected genes have been widely applied in clinics or research to track changes in the biological systems of cells. For example, the activation of IL7R could initiate precursor B-cell acute lymphoblastic leukemia [33], KLRB1 showed a suppression in human cancer tissues [34], and NKG7 regulated cytotoxic granule exocytosis and inflammation [35]. Overall, our experiments demonstrated that scMinerva’s predictions could be used to detect biomarkers in both coarse-grained and fine-grained cell classes. The detection of these biomarkers could contribute to a better understanding of biological processes and disease mechanisms, potentially leading to the development of new diagnostic tools and treatments for various diseases.

Based on the results of our biomarker detection experiments, we sought to interpret our model by considering the following intuition: during the transition and walk generation, nodes with a reasonably high feature expression level serve as “Key Opinion Leaders” in the graph network. These nodes are often identified as anchor nodes in node classification tasks, such as in the KNN algorithm. As such, we were curious about their role during the random walk, particularly for sparse single-cell data that lacks high library-size cells.

To test our intuition that nodes with high feature expression levels act as “Key Opinion Leaders” in the graph network during the transition and walk generation, we computed nodes’ occurrence frequency in these walks and checked the expression levels of marker genes detected in Figure 5a-c. Due to some genes being absent from the raw data, we only included the reserved genes in the figure. Our analysis revealed that, after training, the generated walks and output embeddings were more heavily influenced by nodes with high feature expression levels. Figure 5**d** shows the expression level of the highest occurrence frequency of 20 nodes before and after training. It can be observed that the gene expression levels of the top 20 nodes before training had no significant differences. However, after training, we found that the top 20 nodes mostly exhibited high gene expression levels. This comparison strongly suggests that after training with GCN, nodes with a higher occurrence frequency are more likely to upregulate the marker genes than low occurrence frequency nodes. If we assume that nodes with a higher chance of being walked have a higher priority, then GCN assigns a higher priority to high expression level nodes and utilizes information from their neighbors more frequently. These nodes are particularly important in sparse single-cell data as they retain more valid information about their cell type. In other words, our method broadens its knowledge space from these more valuable cells in single-cell data, making its output more reasonable as it benefits strongly from the representatives of different cell types in the graph network. This discovery establishes a connection between our framework and this biological problem.

Understanding the effect of these highly expressed nodes is essential for gaining insights into the underlying biological processes and for developing more effective algorithms for analyzing single-cell data. ScMinerva highlights the importance of considering the role of highly expressed nodes in the analysis of single-cell data. This, in turn, can lead to the development of more effective treatments and diagnostic tools.

### 2.6 scMinerva reveals potential differentiation changes of naive immune cells after infection of COVID-19

Since scMinerva has shown its ability to provide biomarkers correctly and the gene activity value on biomarkers can expressively reveal the cell stage in a fine-grained manner, we continued to use the predicted results of scMinerva to analyze the cell differentiation trend at the single-cell level. In this section, we use the predictions from scMinerva to compare the differences in cell differentiation between SARS-CoV-2 (COVID-19) infected and healthy human blood immune cells from the COVID-PBMC dataset [24]. We use the MELD [36] method to analyze the potential differentiation trend of Naive T cells in patients. The selected infected cells are from critical symptoms of human beings and can reflect the changes in the long term. T cells play an important role in human immunity, however, most existing analyses on the impact of COVID-19 on T cells are at a cluster level. Researchers infer the impact from changes in cell-type proportions observed in different symptom durations. In our study, we analyze the potential differentiation tendency of Naive CD4 T cells at a single-cell resolution using MELD and conclude results that confirm some recently proposed hypotheses at a single-cell level. To avoid repetition, we only analyze CD4^+^ T cells in this section and observe the potential changes in cell differentiation after infection.

To begin our analysis, we first examine the healthy cells in the COVID-PBMC dataset using MELD, which takes the raw expression data and predicted cell labels from scMinerva as input. MELD generates a sample density for each cell in relation to different cell types. This “sample density” is a kernel density estimate that represents the likelihood of the sample label given the data. The resulting output has rows representing cells and columns representing cell types. Each entry in the sample density reflects the kernel density estimate for a cell on a specific cell type. In our case, we input 10 unique cell types in the label. In essence, the cell type that a cell obtains the highest sample density is the most likely type that the cell will differentiate into.

Subsequently, we focused on CD4^+^ Naive cells and applied Gaussian Mixture Model (GMM) with 10 components to estimate their density on the CD4^+^ Naive column. As illustrated in Figure 6**a**, the majority of cells had high sample density on CD4^+^ Naive, with only a small fraction falling below 0.05. To validate the predictions from scMinerva, we replaced the input labels with the predicted labels and repeated the same analysis. Figure6**b** shows that the predictions from scMinerva corresponded well with the annotations. In Figure 6**c**, we visualized the sample density scores on important functional CD4 T cell types based on the predictions from scMinerva. For healthy cells, there was no significant differentiation tendency in most cell types, with the sample density from different cell types being relatively uniform (see Figure 6**c**). We provide a complete graph for all 10 cell types in appendix.

**Figure 6:**
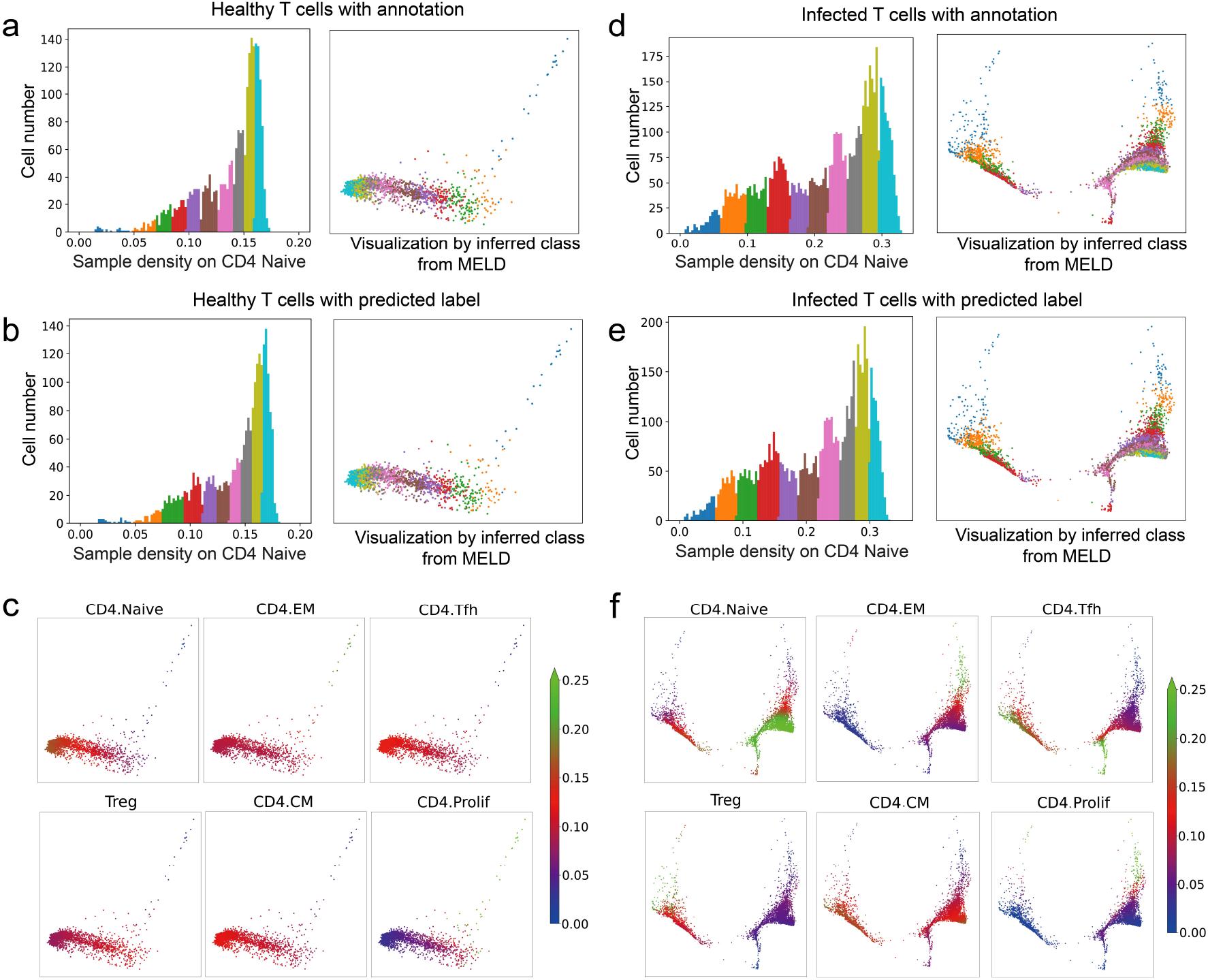
Cell differentiation analysis on CD4 Naive T cells on healthy samples and COVID-19 infected samples. **a**. The clusters fitted by Gaussian Mixture Model(GMM) on healthy tissues inferred by annotations. **b**. Same as **a**, but is inferred by predictions of scMinerva. Our method shows a strong approximation to the results of annotations. **c**. Differentiation likelihood interred by predictions. The differentiation is not active on nearly all cell types. **d**. The clusters fitted by GMM on infected cells inferred by annotations. **e**. Same as **d**, but is inferred by predictions of scMinerva. Our method shows a strong approximation to the annotations. f. Differentiation likelihood inferred by predictions. The differentiation to CD4^+^ T_CM_, CD4^+^ T_FH_, CD4^+^ prolif are activated.

Then, we repeat the procedure on COVID-19-infected cells. Similarly, Figure 6**d,e** compare the GMM modeled sample density on CD4^+^ Naive column by ground-truth label and predictions of scMinerva respectively. Our prediction also shows a great approximation to the annotations. Following this, we visualized the differentiation tendency on some important functioning CD4 cell types in Figure 6**f**. It can be observed that infected cells are under a prosperous differentiation in comparison to healthy cells, especially to CD4^+^ prolif, CD4^+^ T_FH_, CD4^+^ T_EM_, CD4^+^ T_CM_ and CD4^+^ T_reg_. We provide a full graph concerning all ten classes in appendix.

Our results are consistent with the findings of Jung J.H. *et.al.* [37], who reported a significant number of cells differentiating into diverse memory subsets, including CD4^+^ T_EM_ and CD4^+^ T_CM_, in COVID-19 infected tissue compared to healthy tissue. However, their conclusions were based on observations at the bulk level, whereas our study confirms this phenomenon at the single-cell level, specifically in under-differentiated CD4^+^ Naive cells, as shown in Figure 6**f**. The proliferation is also supported by other functioning T cells, such as CD4^+^ T_H_1, CD4^+^ T_H_2, and CD4^+^ T_FH_, as demonstrated in the full graph in appendix. We observed cell proliferation in COVID-19 infected cells at various stages of symptom duration, with a more pronounced effect in critical patients due to T cell apoptosis [38]. These results reflect changes in the biological system and have important practical implications for clinical research.

## 3 Discussion

We have developed scMinerva, an unsupervised method for integrated analysis of single-cell multi-omics datasets. Our method possesses key features that distinguish it from previous methods. Firstly, scMin-erva is more practical to adapt to annotation versions with different granularity. This is achieved through an independent fine-tuning process that effectively classifies cell-types or cell-stages. In contrast, previous unsupervised methods typically complete the cell-type annotation task by clustering the cells first and then using the aggregated cluster-level expression profiles and marker genes to label each cluster. However, the suitability of the fine-tuning process for scMinerva makes it more powerful and accurate for cases requiring fine granularity annotation. Secondly, scMinerva’s embeddings powerfully reserve global sample (dis)similarities and correlations, which contributes to its overall excellent performance on singlecell integrated classification even with a classifier fine-tuned with only 5% labels. This demonstrates the label efficiency of scMinerva. Thirdly, scMinerva is robust to noise and has the ability to capture valid biological information from different omics data. It performs stably on noisy datasets, while other methods are strongly impaired by low-quality omics data.

Our method formulates the integrated analysis task as a heterogeneous graph learning problem and proposes a novel algorithm omics2vec. This algorithm works jointly with a GCN model and enables a more comprehensive inference. The analysis of walks generated by omics2vec combined with the knowledge of biomarkers provides an interpretable process to understand how our method efficiently captures complementary information from different omics data. Interestingly, we use the results from scMinerva to identify meaningful biomarkers and analyze single-cell differentiation trends. Our findings are consistent with clinical discoveries and reveal significant potential for biomedicine research.

While scMinerva has demonstrated excellent integrative classification performance in many scenarios, we have observed that our method performs poorly when handling datasets measured by the scNMT technique [26]. Interestingly, in the benchmarking process (Figure 2b), we have found that other tools’ performance also drops when handling scNMT datasets. However, we have evaluated some fully-supervised methods on this dataset and found that some of them perform well. This phenomenon suggests that the current unsupervised or weakly-supervised methods might not solve this problem effectively and that further research is needed to address this issue. Regarding clinical data prediction, scMinerva has proven to be capable of stably predicting popular measurement techniques (i.e., CITE-seq, ASAP-seq, etc.) in clinical cases with statistical power. Our results are consistent with previous related clinical studies [37, 38].

In conclusion, scMinerva is a superior method for the integrated analysis of single-cell multi-omics datasets, with several key advantages over previous methods. Our approach is more powerful for accurate and fine-grained annotation, achieves label efficiency, and demonstrates robustness to noise and ability to capture valid biological information from different omics. We have formulated this analysis task as a heterogeneous graph learning problem and proposed a novel algorithm, omics2vec, which, when combined with a GCN model, enables a more comprehensive inference. Our results show that scMinerva outperforms previous methods in terms of flexibility and simplicity, and is capable of integrating any number of omics types. Furthermore, our method is versatile and has been effectively applied to biomarker detection and analysis of single-cell differentiation trends, making it a valuable tool for researchers in various fields. With its ability to seamlessly integrate with other tools, scMinerva has the potential to facilitate exploration of the connection between single-cell data and numerous clinical discoveries.

## 4 Method

### 4.1 Datasets, annotations and pre-processing

In this section, we provide detailed information on the data preprocessing and the cell-type information for each dataset used in our analysis.

#### GSE128639

The multi-omics data matrices were used as quantified in the original experiments [23]. For gene expression, standard log-normalization with default parameters in Seurat [23] was conducted. The only difference with the original implementation in the paper is that we take the raw data of HTO separately from the dataset as the third omics. HTO is extremely sparse data so with this as a third omics, the performance of Seurat 4.0 will be strongly lagged back. The cell-type information took into consideration of both known RNA and protein markers. They placed clusters into eight broad groups and further subdivided these groups into 30 level 2 annotation categories. For the details, please refer to the appendix of the reference [23].

#### GSE156478

For the GSE156478 dataset, we followed the same data and cell-type information processing as presented in scJoint [39]. To refer to the detail, please check its data preprocessing section in the appendix. Briefly, the control and stimulated CITE-seq were filtered based on the following criteria: mitochondrial reads greater than 10%; the number of expressed genes less than 500; the total number of UMI less than 1000; the total number of ADTs from the rat isotype control greater than 55 and 65 in the control and stimulated conditions respectively; the total number of UMI greater than 12,000 and 20,000 for the control and stimulated conditions respectively; the total number of ADTs less than 10,000 and 30,000 for control and stimulated conditions respectively. The cells that were classified as doublets in the original study were filtered out. For the ASAP-seq data, cells with a number ADTs more than 10,000 and number of peaks more than 100,000 were filtered out. Finally, 4502 cells (control) and 5468 cells (stimulated) from ASAP-seq, 4644 cells (control), and 3474 cells (stimulated) from CITE-seq were included in the downstream analysis. The number of common genes across the four matrices is 17441 and the number of common ADTs is 227 [39].

#### scNMT

The multi-omics data and cell-type information were obtained from the original study [40]. Generally speaking, gene counts were quantified from the mapped reads by featureCounts [41], and gene annotations were pbtained from Ensembl version 87 [42]. Only protein-coding genes mathcing canonical chromosomes were considered. For methylation and accessibility pseudo-bulk profiles, the values were averaged using running windows of 50 bp. The information from multiple cells was combined by calculating the mean and the standard deviation for each running window. Accessibility profiles were processed with each cell and gene in +/- 200 bp windows around the TSS. Only genes covered in at least 40% of the cells with a minimum coverage of 10 GpC sites were considered [26].

#### SNARE

SNAREseq [25] consists of chromatin accessibility and gene expression. The data is collected from a mixture of human cell lines: BJ, H1, K562, and GM12878. We reduce the dimension of the data by PCA. The size of the resulting matrix for scATAC-seq is of 1047 × 1000 and 1047 × 500 for the gene matrix. We use the code provided by the author to generate annotations for BJ, H1, K562, and GM12878. The cell-type information was obtained from the original study [25].

#### COVID-PBMC

The data and cell-type information were obtained from the original study [43]. Briefly, FASTQ files were generated from raw sequencing reads by the Cell Ranger mkfastq pipeline. Cell Ranger count pipeline (v3.1) was utilized to perform alignment, filtering. barcode counting, and UNI counting. GRCh38 was denoted as genome reference. To remove dead and dying cells, Cells with mitochondrial gene percentages higher than 12% and cells with less than 200 genes was filtered out. For CITE-seq samples, the cells were demultiplexed and hashing adt COUNTS were removed. The remaining counts were normalized by library size and square. For TCR data, the raw sequencing reads of the T cell receptor (TCR) libraries were prcessed by the Cell Ranger V(D)J pipeline by 10x Genomics. Only V(D)J contigs with high confidence defined by cell ranger were considered. The cells of one beta chain contig and zero or one alpha chain contig were remained [43].

### 4.2 Framework of scMinerva

#### 4.2.1 Data simulation from real-world datasets

We generate a four-omics dataset based on the synthetic RNA-seq generated by Splatter [21]. To maintain the mapping between different modalities, we train three Feedforward Neural Networks (FNN) to simulate the mapping between different modalities, including mappings from scATAC-seq to scRNA-seq, from scRNA-seq to ADT matriX, and from scATAC-seq to ADT matrix. We utilize the real-world datasets mentioned below to train these three GNNs. Firstly, we create synthetic scRNA-seq by Splatter, the dimension of which is equal to scRNA-seq from sci-CAR [3]. Then we map the generated scRNA-seq to scATAC-seq by the FNN we trained upon sci-CAR. Similarly, we generate two other ADT matrices from simulated scRNA-seq and scATAC-seq. These two models are trained with scRNA-seq, scATAC-seq, and ADT matrices from GSE156478 and we utilize PCA to make the dimension of scRNA-seq and scATAC-seq consistent with sci-CAR so that we can generate a set of data. We developed four sets of data, with five classes and sample numbers 2k, 5k, 10k, and 30k, respectively. The simulated RNA, ATAC, ADT from RNA, and ADT from ATAC data are of feature numbers 815, 2613, 227, and 227 respectively.

#### 4.2.2 Preliminaries

Consider a dataset with *c* omics and *n* samples (*i.e.*, cells). Denote the samples set *S* = {*s_i_*}, *i* ∈ [1,*n*]. We consider the sample cells as nodes, the gene activity value of sample cells in the omics as node features and the similarities (*i.e.*, Euclidean distance) between nodes pairwisely as their edge weights. The biological problem is formulated in such a graph setting. Throughout the paper, 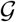 stands for a graph, and 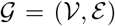 where 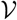 refers to its node set and 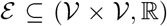 is its edge list. 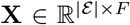 stands for the feature matrix where *F* is the amount of features and a node embedding **x**_*i*_ is a row of **X** of dimension ℝ^*F*^. The adjacency matrix of 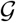 is defined as 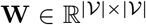 where **w***_v_a_v_b__* stands for the weight of the connecting edge from node *v_a_* to node *v_b_*.

Our objective is to learn a unified embedding for each sample in *S* integrating all the omics. To learn the embedding, We will first construct a heterogeneous graph 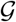 from a list of sub-graphs 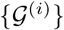. Following the above scheme, we define the corresponding subgraph for the *j*-th omics as 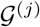 and define 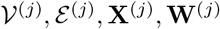 accordingly. For the heterogeneous graph, we define it as 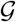 and define 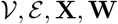 accordingly.

**Algorithm 1.**
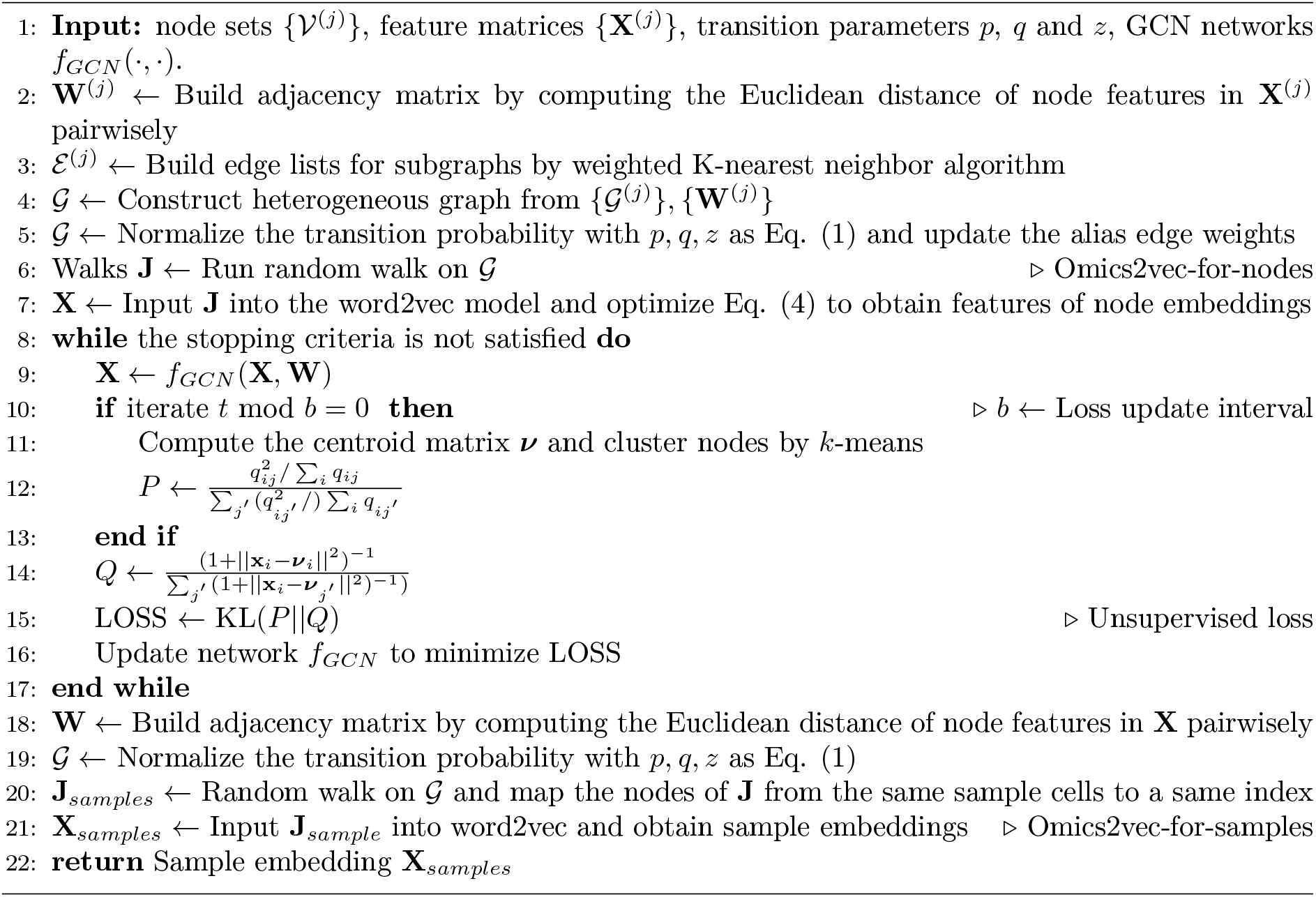
Pseudo-code of scMinerva

#### 4.2.3 Heterogeneous graph construction

The problem is formulated as a graph learning problem and to begin, a heterogeneous graph 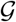 is initialized as the input. Given the feature matrices {***X***^(*j*)^} and node sets 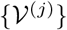 for each omics, the Euclidean distances between the nodes are calculated pairwisely to form {**W**^(*j*)^ | **W**^(*j*)^ ∈ ℝ^*n*×*n*^}. The K-nearest neighbor algorithm is then used to construct the edge lists 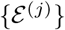 and define the list of subgraphs 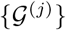.

To aid further discussion, the term **o**(*v*) is defined as the *omics index function* of node *v* and *v_rj_* represents the mapping node of sample *s_r_* in the *j*-th omics, *i.e., **o**(*v_rj_*) = *j*.*

The heterogeneous graph 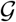 is then constructed, containing all nodes and edges from 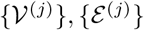. To facilitate inter-omics transitions, initial values of the omics-to-omics edges are assigned to be the unity. Formally speaking, the adjacency matrices {**W**^(*j*)^} are appended diagonally to form **W** ∈ ℝ^*nc*×*nc*^ and nodes from the same sample in different omics are linked with an edge weight of 1. Denote **w**_*ab*_ as the edge weight between node *a* and node *b* and its value is assigned by the (*a*-th, *b*-th) entry of matrix **W**. Formally, **w***_v_rt_v_rl__* = 1 for all *v_rt_* and *v_rl_* where *t* ≠ 1. Undefined entries in **W** are assigned a small enough constant, *ϵ* = 1*e* – 4, to prevent zero division. The new graph has 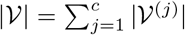 nodes and 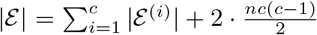 edges, with 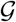 being a directed graph, particularly for the omics-to-omics edges (note the coefficient of 2). In the next steps, the adjacency matrix **W** and the feature matrix **X** of 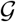 will be optimized.

#### 4.2.4 Omics2vec

Now we will introduce our *omics2vec* algorithm, which is inspired by word2vec [17] and node2vec [18]. It fits node2vec to this biological problem and enables integrated analysis on the heterogeneous graph built on multi-omics data. Briefly, word2vec is a model that learns to embed words in a high-dimensional space where semantically similar words are close to each other [17]. Furthermore, node2vec extends the idea of word2vec to graph-structured data [18]. It learns to embed nodes by defining a biased random walk procedure that explores the graph nodes’ similarity in a way that balances the exploration of local and global network structures. By viewing different nodes in walks as words, node2vec inputs the generated walks into word2vec and ensures the embeddings of spatially close nodes are similar to each other.

Node2vec is a graph search algorithm that combines Breadth-First Search (BFS) and Depth-First-Search (DFS) techniques. It introduces two hyper-parameters, *p* and *q*, to control the second-order random walk. The parameter *p* controls the probability of revisiting a node, while *q* controls the in-out forward (for the detailed geometric meaning of these parameters, please refer to Eqt. (2)). We further investigate the parameter sensitivity of this algorithm, please refer to the appendix. By multiplying the multiplication inverses of these hyper-parameters with the edge weights and normalizing the transition probability to 1, node2vec sets up a biased random walk procedure. However, applying node2vec to this biological heterogeneous graph presents a challenge because the inter-omics transition edges are undefined. Moreover, the weights of the inter-omics edges play a significant role in the performance, as they determine the probability of being explored in the neighborhoods of a sample’s counterparts on different omics. To overcome this challenge, we propose a novel algorithm called omics2vec, which extends the node2vec idea to analyze biological heterogeneous graphs. Omics2vec enables inter-omics transition and can work together with the GCN model training (Section 4.2.5), which optimizes interomics edge weights. To compute the transition probability on graph 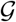, we combine the edge weights **W** and transition-control hyper-parameters, generate random walks based on these probabilities, and input them into the word2vec model to generate node embeddings. Omics2vec has two types: embed for nodes and embed for the samples (single cells). The two types share the same first few steps as the following illustration:

To represent the transition probability from one node *b* to another node *a*, a transition probability function *P*(*a*|*b*) is defined. To generate random walks of a fixed length *l* in graph 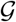, we denote *α_i_* as the *i*-th step in the walk *α* starting from *α*_0_. Steps *α_g_* | *g* ∈ [1, *l*] are generated by the following probability function:

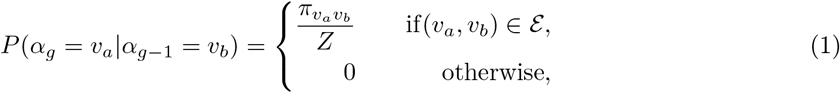

where *π_v_a_v_b__* is the unnormalized transition probability from node *v_b_* to node *v_a_* that is defined in the following contents and *Z* is a normalizing constant.

The normalizing constant *Z* is carefully selected to guide the exploration of different types of neighborhoods. Instead of the mixture of BFS and DFS used by node2vec, we guide the biased random walk with three parameters *p, q*, and *z* all of whom are positive values. The parameter *z* is a new addition to the algorithm and can be seen as the “omics-first search”.

Considering a random walk that just traveled from node *v*_1_ (i.e., step *α*_*g*–1_ = *v*_1_) to node *v*_2_ (i.e., step *α_g_* = *v*_2_), the walk now is determining the next step *α*_*g*+1_. So it evaluates the transition probability *π*_*v*_2_*v*_3__ on edge (*v*_2_, *v*_3_) leading from *v*_2_. We set the unnormalized transition probability as *π*_*v*_2_*v*_3__ = *α_pqz_* · **w**_*v*_2_*v*_3__. Here, *α_pqz_* is a function that depends on the hyperparameters *p, q*, and *z*, and the *shortest distance d*_*v*_1_*v*_3__ between nodes *v*_1_ and *v*_3_ in the graph:

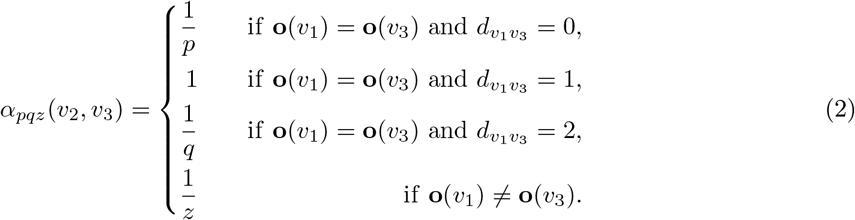

Specifically, if nodes *v*_1_ and *v*_3_ belong to the same omics type and have a shorter distance of 0, then 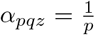. If *p* is smaller, it means that the algorithm has a higher probability of revisiting the node in the last step. If the distance between *v*_1_ and *v*_3_ is 1, then *α_pqz_* = 1, indicating that the algorithm can explore nodes in the same omics type that are immediate neighbors. If the distance between *v*_1_ and *v*_3_ is 2, then 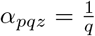. This setting allows the algorithm to explore nodes in the same omics type that are not immediate neighbors. Finally, if *v*_1_ and *v*_3_ belong to different omics types, then 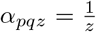. This setting allows the algorithm to explore nodes and moreover, their neighborhood, in other omics types. Once the transition probabilities are calculated, the algorithm generates walks randomly, starting from each source node and repeating the random walk *r* times for each source node. The number *r* is a hyperparameter that determines the number of walks generated from each source node.

For the next few steps, Omics2vec-for-Nodes and Omics2vec-for-Samples are different. Omics2vec-for-Nodes will directly input the generated walks into the word2vec model which outputs the node embeddings. However, omcis2vec for samples will conduct an additional index mapping step for generated walks which maps the node indices to their sample indices. After the mapping, we obtain the walks that reflect the sample’s similarities in the perspective of the global heterogeneous graph. By inputting the walks for samples after mapping, we obtain the embeddings for sample cells. Next, we will elaborate on these steps in more detail.

##### Omics2vec-for-nodes

When the random walk finishes, these are 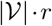 walks in total. Since each walk is of length *l*, the walk matrix 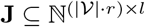 is the input of the word2vec model. Let *f*: *S* → ℝ^*F*^ be the mapping function from the node set to embeddings of dimension *F*. We aim to learn such representations for the later tasks. For source node, 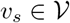, define 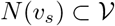 are the neighborhood of node *v_s_* generated through a sample of walks. With a skip-graph architecture, we are trying to find the *f* that gives

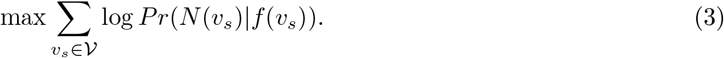

The above objective function maximizes the log-probability of observing *N*(*v_s_*) for node *v_s_* conditioned on its mapping after function *f*. Here, by assuming conditional independence among observing different neighborhood nodes given the feature representation of the source, Eq. (3) can be simplified to:

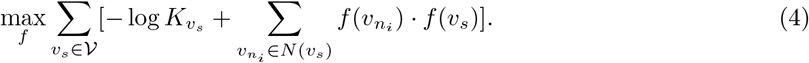

where 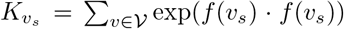 is the partition function for nodes. Solve Eq. (4) using stochastic gradient ascent over the model defining the features *f*. The output embeddings from *f* is the feature matrix 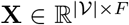 of nodes on the heterogeneous graph 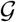. After obtaining the feature matrix **X**, we can improve it using graph convolutional networks (GCNs). GCNs are a type of neural network that operates on graph-structured data and can learn representations by aggregating information from the node’s local neighborhood. By applying GCNs to the feature matrix **X**, we can learn more expressive node representations that take into account the structure of the heterogeneous graph.

##### Omics2vec-for-samples

It is similar to the Omics2vec-for-Nodes algorithm. However, in the final step of our method, we use the walks generated by omics2vec to map the indices of nodes from different omics to their indices of sample cells. Specifically, we ensure that the counterparts of one sample cell from different omics are mapped to the same sample cell in the walks. This remapping produces a new set of walks, denoted as **J**_sample_, which contain information about the relationships between sample cells across multiple omics. Next, we input the remapped walks **J**_sample_ into the word2vec model to obtain the embedding of sample cells **X**_sample_. This embedding captures the integrated multi-omics information by leveraging the well-clustered neighborhood information from the entire heterogeneous graph.

#### 4.2.5 Model training

With the node features **X** as well as the graph adjacency matrix **W** as inputs, we train a two-layer GCN model *f_GCN_*(·, ·) to optimize the graph. To train the GCN, we adopt an unsupervised clustering loss function DeepCluster [44] which produces pseudo-labels and learns from the iteratively updated cluster assignments. The training enforces the nodes with higher similarity to be closer in the embedding space and the nodes with low similarity to be dispersed.

To apply this loss, we first run *k*-means algorithms to cluster the output feature matrix **X** into *k* different groups based on the geometric neighborhood. *K*-means outputs a set of optimal pseudolabels and we denote it as {*y*\*}. Our loss supervising GCN is based on minimizing the Kullback-Leibler (KL) divergence between a student’s t-distribution kernel *Q* to the clusters and a target distribution *P*. The target distribution *P* is a Gaussian distribution centered at each cluster center. The student’s t-distribution kernel *Q* is used to calculate the soft assignment probability *q_ij_* of the embedding **x**_*i*_ to the cluster centroid *v_i_*, 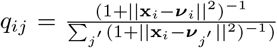. Next, based on *q_ij_*, a target distribution *P* is calculated to help learn from the assignments with higher scores where 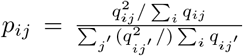. Finally, the loss function is defined as

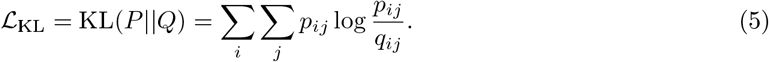

We optimize the feature matrix **X** by minimizing Equation 5. We continue to iterate until the loss is small enough (*i.e.*, 5e-2), at which point we terminate the iteration and output the embeddings **X** for nodes in 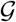. By optimizing the feature matrix, we ensure that similar nodes from the heterogeneous graph are gathered together in the embedding space. Once we have the node embeddings, we compute the distance of nodes pairwise and reconstruct the heterogeneous graph from these embeddings. The omics-transition edges are updated by the results after GCN training, resulting in more reasonable edges than the naive initialization (assigning a unity value to all the inter-omics edges). These updated edges can improve downstream performance as shown in the ablation study (Section 2.3). Then, we run Omics2vec-for-Samples to generate walks following the process in Section 4.2.4 and obtain the sample embeddings. Then we will evaluate the model through multiple experiments.

#### 4.2.6 Experiment settings

##### Cell classification

Previous methods of cell classification tasks mostly are conducted in a fully-supervised manner. As we have mentioned in the introduction, they could not quickly be adapted to fit different grain-level annotations. Jointly considering the labor-intensive and time-consuming processing of data annotation, we evaluate the label-efficiency of methods with a simple nearest-neighbor classifier which is independent of the unsupervised training stage. In detail, for the produced embeddings of scMinerva and other methods, we fit them with independent nearest neighbor-based classifiers and fine-tune them using only 10% annotations of the whole training set.

We implement existing state-of-the-art methods with similar approaches as illustrated in our introduction, including DeepMAPS, CiteFuse, totalVI, and Seurat 4.0 (weighted nearest neighbor), in their recommended settings. For MOFA+ and Conos, the dimension of their embeddings needs to be enlarged to fit a large dataset. So we search the embedding dimension of {100, 200, 300, 400} and select the best-performing dimension. Notably, MOFA+ is very memory-consuming, and we failed to conduct the experiment with the originally preprocessed data using a RAM of 32GB when the number of samples is greater than 10k. Thus, we perform PCA to reduce the input dimension, where we also search among {100, 200, 400, 800} to select the optimal performance. Besides, DeepMAPS, CiteFuse, and TotalVI cannot process three-omics data, and therefore they are excluded in the chart for COVID-PBMC, scNMT, and GSE128639. To evaluate the quality of the generated embeddings, we perform classification by fitting a K-nearest Neighbor (KNN) Classifier with the number of neighbors as 30 for datasets containing more than 5k samples, and with the number of neighbors as 8 for datasets smaller than 5k.

##### Cell differentiation analysis

Cell differentiation is the process by which a single cell develops into many different specialized cell types with unique functions. It is important for understanding cell development, disease, and drug and for developing new therapies and treatments. In this paper, we take cell differentiation analysis as an example to show the practice value of our model together with MELD [31]. We take human blood immune cells from dataset COVID-PBMC [45] to compare the differences in cell differentiation between cells infected with SARS-CoV-2 (COVID-19) and healthy cells. To avoid repeating, we only take *CD4*^+^ T cells which contain 10 sub-classes to observe the potential cell differentiation changes. Since there are 10 sub-classes, we run the Gaussian Mixture Model (GMM) with the number of components as 10 and all other parameters are the same as the default setting in MELD.

### 4.3 Hyperparameters

#### Omics2vec

We run random walk on the heterogeneous graph with three transition controlling parameters named *p, q*, and *z*, where *p* controls the likelihood of immediately revisiting a node in the walk, *q* allows the search to differentiate between “inward” and “outward” nodes, and *z* controls an inter-omics transition within the frame. By default, we set *p, q, z* all equal to the unity. To ensure a rich connection between omics and avoid zero division during normalizing, we also introduce a hyper-parameter *δ* to smooth the graph. In another word, the omics-transition links will have a small enough default value equal to *δ* before normalizing. We set *δ* to 1*e* – 4.

For other random walk parameters, we follow the default value set in node2vec [18]. In detail, on each node of the graph, we run random walk start from it 10 times. Each time it will generate a walk of length 80. With the generated walks, we input them to word2vec under algorithm CBOW and window size 10. Word2vec will output embeddings of dimension 128 for nodes contained in walks.

#### GCN model

The GCN architecture has two primary hyperparameters: the latent dimensions of the autoencoders and the *k* of k-means that DEC clusters the nodes. We simply set *k* = *c* × #cell-type (number of cell-types) and stop the training while the loss function achieves a value smaller than 5*e* – 2. The dimension of the latent space is set to be 32.

### 4.4 Performance evaluation

#### ACC

We denote *Positive* as *P, Negative* as *N, True positive* as *TP, False negative* as *FN, False positive* as *FP*, and *True negative* as *TN*. Then we can define accuracy (ACC) [46] as

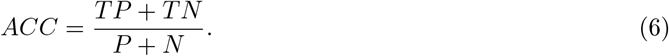

#### F1 score

Here, F1 score can be calculated [47] as:

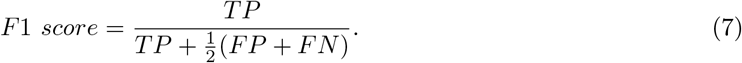

The F1-macro is the arithmetic mean of all the per-class F1 scores, and F1-weighted is computed by taking the mean of all per-class F1 scores considering the weight. Weight refers to the number of actual occurrences of the class in the dataset.

#### ARI

Adjusted rand index (ARI) is used to measure the similarity between the predicted labels and ground truth. The Rand Index (RI) calculates a similarity measure between two clusterings, taking all pairs of samples into consideration. It counts pairs that are assigned in the same or different clusters in the predicted and actual clusterings. ARI is a corrected-for-chance version of the Rand index defined as

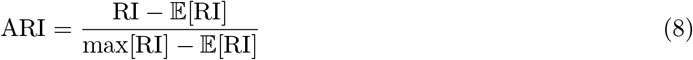

### 4.5 Statistics and reproducibility

Datasets were chosen in order to show the functionality and performance of our method. No data were excluded from the analyses. Replication and randomization are not applicable since we did not collect any experimental data. Hypothesis testing methods are explained in each figure legend. To reproduce the results, please find the Source Data file we provided.

## 5 Funding

This research is funded by the Chinese University of Hong Kong (CUHK) with the award number 4937025, 4937026, 5501517, and 5501329.

Yongshuo Zong was supported by the United Kingdom Research and Innovation (grant EP/S02431X/1), UKRI Centre for Doctoral Training in Biomedical AI at the University of Edinburgh, School of Informatics. For the purpose of open access, the author has applied a creative commons attribution (CC BY) licence to any author-accepted manuscript version arising.

## 6 Data availability

The datasets analyzed in this study are available under the following accession numbers: GSE128639 [23], GSE156478-CITE [22], GSE156478-ASAP [22]), COVID-PBMC [24]), SNARE-seq [25] and scNMT-seq [26].

## 7 Code avalibility

The open-source implementation of scMinerva is available at https://github.com/yistyu/scMinerva, and the experiments conducted to produce the main results of this article are also stored in this repository.

## 8 Ethics approval and consent to participate

Not applicable.

## 9 Competing interests

The authors declare that they have no competing interests.

## 10 Authors’ contributions

T.Y. and Y.L. conceived the project. T.Y. implemented the model. T.Y. and Y.W. generated figures. T.Y., Y.Z., Y.W., and X.W. run the experiments on baseline methods. T.Y. and Y.Z. wrote the manuscript with feedback from all authors. Y.L. supervised the project. The authors read and approved the final manuscript.

## Notes

### Competing Interest Statement

The authors have declared no competing interest.

### Summary of Updates

Improve the presentation of the method part.

